# Heat-treated virus inactivation rate depends strongly on treatment procedure: illustration with SARS-CoV-2

**DOI:** 10.1101/2020.08.10.242206

**Authors:** Amandine Gamble, Robert J. Fischer, Dylan H. Morris, Kwe Claude Yinda, Vincent J. Munster, James O. Lloyd-Smith

## Abstract

Decontamination helps limit environmental transmission of infectious agents. It is required for the safe re-use of contaminated medical, laboratory and personal protective equipment, and for the safe handling of biological samples. Heat treatment is a common decontamination method, notably used for viruses. We show that for liquid specimens (here, solution of SARS-CoV-2 in cell culture medium), virus inactivation rate under heat treatment at 70°C can vary by almost two orders of magnitude depending on the treatment procedure, from a half-life of 0.86 min (95% credible interval: [0.09, 1.77]) in closed vials in a heat block to 37.00 min ([12.65, 869.82]) in uncovered plates in a dry oven. These findings suggest a critical role of evaporation in virus inactivation via dry heat. Placing samples in open or uncovered containers may dramatically reduce the speed and efficacy of heat treatment for virus inactivation. Given these findings, we reviewed the literature temperature-dependent coronavirus stability and found that specimen containers, and whether they are closed, covered, or uncovered, are rarely reported in the scientific literature. Heat-treatment procedures must be fully specified when reporting experimental studies to facilitate result interpretation and reproducibility, and must be carefully considered when developing decontamination guidelines.

**Importance:** Heat is a powerful weapon against most infectious agents. It is widely used for decontamination of medical, laboratory and personal protective equipment, and for biological samples. There are many methods of heat treatment, and methodological details can affect speed and efficacy of decontamination. We applied four different heat-treatment procedures to liquid specimens containing SARS-CoV-2. Our results show that the container used to store specimens during decontamination can substantially affect inactivation rate: for a given initial level of contamination, decontamination time can vary from a few minutes in closed vials to several hours in uncovered plates. Reviewing the literature, we found that container choices and heat treatment methods are only rarely reported explicitly in methods sections. Our study shows that careful consideration of heat-treatment procedure — in particular the choice of specimen container, and whether it is covered — can make results more consistent across studies, improve decontamination practice, and provide insight into the mechanisms of virus inactivation.

## Introduction

The COVID-19 pandemic has led to millions of infections worldwide via multiple modes of transmission. Transmission is thought to occur via respiratory particles expelled by individuals infected by the causative virus, SARS-CoV-2 [1–3]. Epidemiological investigations that environmental transmission of SARS-CoV-2 occurs [4]; this is possible because the virus remains stable for a period of time on inert surfaces and in aerosols [5, 6]. Environmental transmission has been suspected or demonstrated for many other viruses, including hepatitis viruses [7], noroviruses [8], and influenza viruses [9] among others. Rapid and effective decontamination methods can help limit environmental transmission during infectious disease outbreaks. Heat treatment is a widely-used decontamination method, notably used for viruses [10]. It is thought to inactivate viruses principally by denaturing the secondary structures of proteins and other molecules, resulting in impaired molecular function [11]. Heat is used to decontaminate various materials, such as personal protective equipment (PPE), examination and surgical tools, culture and transportation media, and biological samples [12–15]. The United States Centers for Disease Control and Prevention recommends moist heat as a SARS-CoV-2 inactivation method [16].

In this context, multiple studies have evaluated the effectiveness of heat to inactivate coronaviruses on various household surfaces, PPE, culture and transportation media, and blood products [14, 17–22]. Heat-based decontamination procedures are also used for many other viruses, including hepatitis viruses [23], influenza viruses [24], parvoviruses [25], and human immunodeficiency viruses [26].

There are multiple ways to apply heat treatment. Heat can be dry or moist. Heating implements can differ in degree of heat transfer: for example, heat blocks in theory allow more efficient heat transfer than ovens, so samples should more rapidly reach and better maintain the target temperature. Different levels of evaporation may be permitted: for example, samples deposited on flat surfaces or contained in open plates will evaporate more than those in closed vials; both types of container are commonly-used. Local temperature and humidity impact virus inactivation rates by affecting molecular interactions and solute concentration [27]. It follows that factors such as heat transfer and evaporation, which determine solute concentration and alter micro-environment temperature through evaporative cooling, could modulate virus inactivation rates just as ambient temperature does.

We assessed the impact of heat-treatment procedure on SARS-CoV-2 inactivation. We studied dry heat treatment applied to a liquid specimen (virus suspension in cell culture medium), keeping temperature constant (at 70°C) but allowing different degrees of heat transfer (using a dry oven or a heat block) and evaporation (placing samples in closed vials, covered plates or uncovered plates). We then compared the half-lives of SARS-CoV-2 under these different procedures. In light of our results, we reviewed the literature to assess whether heat-treatment procedure descriptions are detailed enough to allow replication and inter-study comparison. We focused our literature review on coronavirus inactivation.

## Results and Discussion

### Estimation of SARS-CoV-2 half-life under four distinct heat-treatment procedures

We prepared a solution of cell culture medium containing SARS-CoV-2, and exposed it to 70°C heat using four different procedures: (1) an uncovered 24-well plate, (2) a covered 24-well plate (using an unsealed plastic lid), (3) a set of closed 2 mL vials in a dry oven, and (4) a set of closed 2 mL vials in a heat block containing water (Fig. 1A). The inactivation rate of SARS-CoV-2 differed sharply across procedures. There were large differences in the time until the virus dropped below detectable levels, despite comparable initial quantities of virus (estimated mean initial titer ranging from 4.5 [4.1, 5.0] log_10_ TCID_50_/mL for the uncovered plate in an oven to 5.0 [4.7, 5.5] for the closed vials in a heat block, Fig. 1B). We could not detect viable virus in the medium after 30 min of treatment (the earliest time-point) in closed vials heated either in a heat block or in a dry oven; we could not detect viable virus after 90 min in covered plates (Fig. 1B). In uncovered plates, we observed a reduction of viral titer of approximately 1 log_10_ TCID_50_/mL after 60 min. Because macroscopic evaporation was observed in the uncovered plates and was almost complete at 60 min, all the samples were complemented to 1 mL with deionized water at collection. Hence, the slower decrease in viral titer observed in uncovered plates (and, to a lesser extent, in covered plates compared to closed vials) can only be explained by a slower inactivation rate, not by virus concentration due to evaporation.

**Figure 1.**
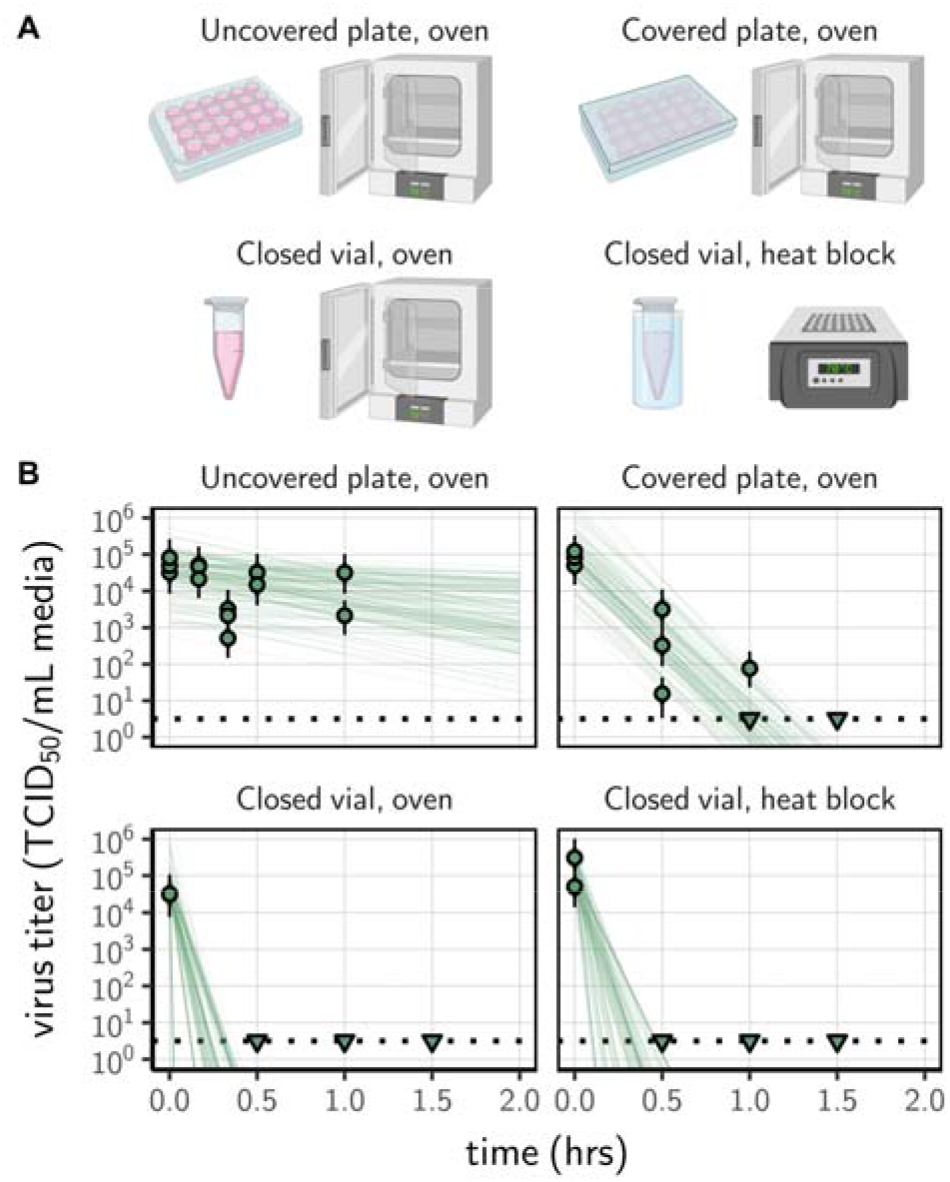
Inactivation of SARS-CoV-2 by heat treatment under different procedures. (A) A solution of SARS-CoV-2 was exposed to 70°C heat. Samples were placed in uncovered or covered 24-well plates, or in closed 2 mL vial before heat treatment using a dry oven or a heat block containing water. (B) Samples were then collected at indicted time-points during heat treatment. Viable virus titer estimated by end-point titration is shown in TCID_50_/mL media on a logarithmic scale. Points show estimated titers for each collected sample; vertical bar shows a 95% credible interval. Time-points with no positive wells for any replicate are plotted as triangles at the approximate single-replicate detection limit of the assay (LOD; denoted by a black dotted line at 10^0.5^ TCID_50_/mL media) to indicate that a range of sub-LOD values are plausible. Lines show predicted decay of virus titer over time (10 random draws per data-point from the joint posterior distribution of the slope and intercept). Panel A created with BioRender.com

Using a Bayesian regression model, we estimated inactivation rates from the experimental data and converted them to half-lives to compare the four procedures. SARS-CoV-2 inactivation in solution was most rapid in closed vials, using either a heat block or a dry oven (half-lives of 0.86 [0.09, 1.77] and 1.91 [0.10, 1.99] min, respectively), compared to the other treatment procedures (Fig. 2; Supplemental Material, Table 1). Inactivation rate was intermediate in covered plates (half-life of 3.94 [3.12, 5.01] min) and considerably slower in uncovered plates (37.04 [12.65, 869.82] min).

**Figure 2.**
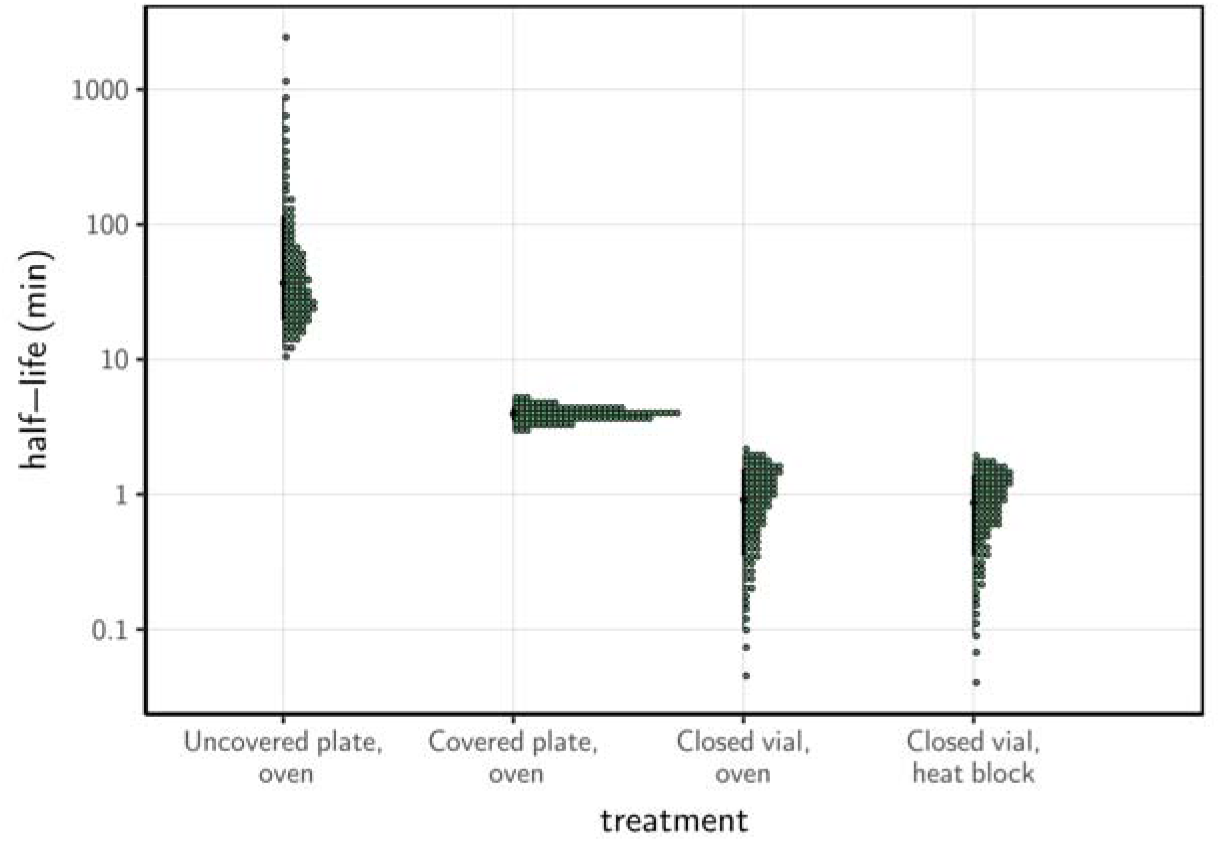
Half-life of SARS-CoV-2 in a solution exposed to 70°C heat under different procedures. Quantile dotplots 69] of the posterior distribution for half-life of viable virus under each different heat-treatment procedure. Half-lives were calculated from the estimated exponential decay rates of virus titer (Fig. 1B) and plotted on a logarithmic scale. For each distribution, the black dot shows the posterior median estimate and the black line shows the 95% credible interval.

**Table 1.** Half-life of SARS-CoV-2 in Dulbecco’s Modified Eagle Medium cell culture medium exposed to 70°C heat under different procedures. Half-lives are calculated from the estimated exponential decay rates of virus titer and reported as posterior median and middle 95% credible interval.

| Procedure | Median (min) | 2.5% | 97.5% |
| --- | --- | --- | --- |
| Uncovered plate, oven | 37.04 | 12.65 | 869.82 |
| Covered plate, oven | 3.94 | 3.12 | 5.01 |
| Closed vial, oven | 0.91 | 0.10 | 1.99 |
| Closed vial, heat block | 0.86 | 0.09 | 1.77 |

The rapid virus inactivation rate seen in closed vials subject to dry heat at 70°C agrees with previously reported results for inactivation of SARS-CoV-2 in virus transportation medium [28], SARS-CoV-1 in human serum [17], and MERS-CoV [18] and canine coronavirus in cell culture medium [21], among other results. All showed a loss of infectivity on the order of 10^4−6^ TCID_50_ after 5–10 min at 65–75°C. None of these studies report sufficient details on their protocol to indicate which of our tested procedures corresponds most closely to their approach. These results have critical implications for real-world heat treatment decontamination practices. Inactivation rates reported in studies that use closed vials may dramatically underestimate the time needed to decontaminate a piece of equipment (uncovered) in a dry oven. We have previously estimated the half-life of SARS-CoV-2 on stainless steel and N95 fabric when exposed to 70°C using a dry oven, without a container to limit evaporation. We found half-lives of approximately 9 and 5 min, respectively [14]. These values are on the same order of magnitude as the half-life of the virus in bulk solution exposed to heat treatment in a covered plate (3.94 [3.12, 5.01] min), and considerably longer than the half-life of the virus exposed to heat treatment in bulk solution in a closed vial. Inactivation rates reported by studies conducted in closed vials should not be used to directly inform decontamination guidelines of pieces of equipment that cannot be treated using the same exact procedure.

### The potential role of evaporation in virus inactivation

The fact that containers that allow more air-flow are associated with slower virus inactivation suggests that evaporation may play a critical role in determining the rate of virus inactivation during dry heat treatment. There are several mechanisms by which evaporation could impact the effectiveness of heat treatment for virus inactivation. First, evaporation could induce a local drop in temperature due to the enthalpy of vaporization of water (or evaporative cooling), limiting the effect of the heat itself. This hypothesis could be verified in future studies by measuring sample temperature (instead of ambient temperature) using a thermocouple. Second, evaporation could lead to modifications of the virion’s solute environment: solutes become more concentrated as the solvent evaporates, and under certain conditions efflorescence (i.e. crystal formation) can occur [29]. Mechanistic modeling of virus inactivation data shows that increased solute concentration increases virus inactivation rate, but efflorescence decreases inactivation rate [27]. Our results show that greater degrees of evaporation during dry heat treatment are associated with slower virus inactivation. This suggests that evaporative cooling, efflorescence, or both may drive lower inactivation rates in closed containers. This could help explain why low ambient humidity levels lead to slow inactivation at high temperatures [30], as low humidity levels allow for more evaporation and possibly efflorescence. The potential role of evaporation as a key modulator of virus inactivation rate is supported by the known importance of other factors that affect evaporation, such as relative humidity [27, 29] and medium composition [20, 31]. We postulate that container shape and surface area to volume ratio will also play a role, as these should also impact the evaporative dynamics. Heat transfer efficiency may also play a role in determining the rate of virus inactivation using dry heat, but our data do not provide evidence for or against this hypothesis, since virus inactivation was extremely rapid in closed vials regardless of whether they were exposed to heat using a dry oven or a heat block.

Our study focuses exclusively on the effect of temperature on virus inactivation. Other factors can affect virus inactivation rate in liquid specimens, for example the composition of the suspension medium [20, 29, 32]. In particular, proteins are thought to have a protective effect on virus viability, while the effect of salts and pH depends on other factors such as ambient humidity [33]. We consider these only implicitly, insofar as they are affected by evaporation. The role of medium composition will be especially important to consider in future studies, as the composition of biological fluids, usually targeted by decontamination procedures, differs greatly from that of cell culture media. In addition, the impact of heat treatment procedure on inactivation rate may differ across microbes. Enveloped and non-enveloped viruses may behave differently from each other, and bacteria may behave differently from viruses [29]. In particular, non-enveloped viruses are generally more stable than enveloped viruses [34], but very few studies have focused on thermal sensitivity [35]. Finally, decontamination procedures must consider not only the effectiveness and speed of pathogen inactivation but also the potential impact of the procedure on the integrity of the decontaminated equipment or specimen. This is particularly important for PPE and for biological samples [14, 36, 37].

Given the substantial effect of heat-treatment procedure on virus inactivation rates, it is critical to specify procedures precisely when comparing inactivation rates between studies or producing guidelines for decontamination. In particular, our results show that protocols that use open containers or uncovered surfaces lead to much slower viral inactivation, at least in bulk medium. For instance, the fact that Chin et al. 2020 [28] used closed vials to quantify SARS-CoV-2 half-life (personal communications) likely gave rise to outliers observed ad 56 and 70°C relative to predicted relationships parameterized from uncovered surfaces [27]. If meta-analyses of the effect of temperature on virus inactivation were to integrate together data collected following different procedures, without corrections, they may lead to false conclusions.

### Reporting of heat-treatment procedures in the literature

Given these findings, we conducted a literature review in order to assess whether heat treatment procedures for coronaviruses were reported with sufficient details to allow reproducibility and appropriate interpretation of results. Our literature review identified 41 studies reporting the effect of temperature on coronavirus stability (Fig. S1), covering 12 coronavirus species and temperature ranging from -70 to 100°C (Table. S1). Among those 41 studies, just 14 included any information about the containers used, and 5 specified whether containers were closed. Only a single study reported container type and container closure explicitly for all experimental conditions [38]. When the information was available, studies of virus stability in bulk liquid medium were always conducted in vials [6, 20, 21, 38–45]. Studies interested in virus stability on surfaces were conducted in vials, in well plates [46] or trays [47], or on surface coupons placed in vials [39] or placed directly on oven rack (personal communication [14]). When specified, vial volume ranged from 1.5 mL to 50 mL [21, 40, 42], and sample volume from 0.001 to 45 mL. Finally, 24 studies included some information about how target temperature (and, in some cases, humidity) conditions were created. Methods included water baths [17, 19, 20, 38, 42, 44, 45, 48, 49], heat blocks [40, 43, 50], incubators [30, 47, 51–54], ovens [14], refrigerators [55–57], isothermal boxes [56], and boxes with saturated salt solutions [58].

This literature review reveals that a variety of setups are used to hold samples and control environmental conditions for virus stability and inactivation experiments. Unfortunately, it also reveals that the vast majority of studies of heat treatment for virus inactivation do not report the exact procedures under which the samples were exposed to heat (in particular whether they were in closed, covered, or uncovered containers). This makes it difficult to compare inactivation rates among studies, and risky to use estimates from the literature to inform decontamination guidelines. More generally, given the potentially large effects of treatment procedure and ambient environment on virus inactivation rate, we recommend that decontamination procedures be validated specifically for the setup to be used, rather than based on inactivation rate estimates from the literature, especially if experimental protocols are unclear.

## Conclusion and Perspectives

Using SARS-CoV-2 as an illustration, we demonstrate that the choice of heat-treatment procedure has a considerable impact on virus inactivation rates in liquid specimens. Our findings highlight the need to better understand the mechanisms controlling inactivation rate, including the role of evaporation. This will require comparative studies including a set of diverse microbes exposed to heat treatment in different conditions likely to impact evaporation dynamics and/or microbe thermal stability, ideally paired with high-resolution physical measurements. These conditions include container sealing, but also sample volume and evaporation surface, medium composition, container material, and heating system. In the meantime, any effort to compare or translate inactivation rates (or even relative patterns) from one setting to another should be undertaken cautiously, accounting for these factors. In particular, as decontamination time can vary by several orders of magnitude across procedures, these factors should be considered when developing decontamination guidelines. Finally, we also call for more thorough description of experimental protocols in scientific publications, for instance through the publication of detailed protocols in online repositories [59], or peer-reviewed journals publishing laboratory protocols. Better understanding the impact of temperature and humidity on virus inactivation is critical not only for designing efficient decontamination protocols but also for predicting virus environmental persistence, with consequences for real-world transmission [27, 60, 61].

## Material and Methods

### Laboratory experiments

We used SARS-CoV-2 strain HCoV-19 nCoV-WA1-2020 (MN985325.1) [62] for all our experiments. We prepared a solution of SARS-CoV-2 in Dulbecco’s Modified Eagle Medium cell culture medium (Sigma-Aldrich, reference D6546) supplemented with 2 nM L-glutamine, 2% fetal bovine serum and 100 units/mL penicillin/streptomycin. For each of the four heat-treatment procedures considered, we placed samples of 1 mL of this solution in plate wells or vials before heat treatment. This relatively low volume was chosen to allow the samples to reach 70°C quickly. The plates were 24-well flat-bottom plates made of crystalline polystyrene with an inner diameter of 15.6 mm (Corning Costar), and 2 mL screw-top vials made of polypropylene with an inner diameter of 10.8 mm diameter (Sarstedt). Both materials have a similar thermal conductivity (0.1-0.13 and 0.1-0.22, respectively, at 23°C) and thickness (about 0.05 mm). The plates and tubes were then placed into either a gene hybridization dry oven or a heat block with water in the wells (Fig. 1A). The large rotating ferris wheel-like apparatus of the gene hybridization oven ensured air mixing during the experiments, preventing a build-up of humid air above the open wells.

Samples were removed at 10, 20, 30 and 60 min from the uncovered 24-well plate, or at 30, 60 and 90 min for the three other procedures. We took a 0 min time-point measurement prior to exposing the specimens to the heat treatment. As evaporation was observed after exposure to heat, all the samples were complemented to 1 mL with deionized water at collection in order to re-hydrate the suspension medium and recover virions with the same efficiency across all treatments. At each collection time-point, samples were transferred into a vial and frozen at -80°C until titration (or directly frozen for experiments conducted in vials). Note that all the samples were kept frozen for 8 days and subject to one freeze-thaw cycle, which may have some (limited) impact on absolute virus titer [63, 64], but not on the estimated inactivation rate (since this depends on relative titers across samples). We performed three replicates for each inactivation procedure. Samples were not exposed to direct sunlight during the experiment.

We quantified viable virus contained in the collected samples by end-point titration as described previously [14]. Briefly, Vero E6 cells were plated the day before carrying out titration. After 24 hours, the cells had reached a confluency of about 85-90% and were inoculated with 10-fold serial dilutions of sample in quadruplicates. One hour after inoculation, inoculum was removed and replaced with 100µL of supplemented DMEM. Six days after inoculation, each well was observed for cytopathogenic effects and classified as infected or non-infected.

### Statistical analyses

We quantified the inactivation rate of SARS-CoV-2 in a solution following different heat-treatment procedures by adapting a Bayesian approach described previously [14]. Briefly, we inferred virus titers from raw endpoint titration well data (infected / non-infected) by modeling well infections as a Poisson single-hit process [65]. Then, we estimated the decay rates of viable virus titer using a regression model. This modeling approach allowed us to account for differences in initial virus titers (0 min time-point) across samples as well as other sources of experimental noise. The model yields posterior distributions for the virus inactivation rate under each of the treatment procedures—that is, estimates of the range of plausible values for each of these parameters given our data, with an estimate of the overall uncertainty [66]. We then calculated half-lives from the estimated inactivation rates. We analyzed data obtained under different treatment procedures separately. We placed weakly informative prior distributions on mean initial virus titers and log virus half-lives. The complete model is detailed in the Supplemental Material.

We estimated virus titers and model parameters by drawing posterior samples using Stan [67], which implements a No-U-Turn Sampler (a form of Markov Chain Monte Carlo), via its R interface RStan. We report estimated titers and model parameters as the median [95% credible interval] of their posterior distribution. We assessed convergence by examining trace plots and confirming sufficient effective sample sizes and *R* values for all parameters. We confirmed appropriateness of prior distributions with prior predictive checks and assessed goodness of fit by plotting regression lines against estimated titers and through posterior predictive checks (SI, Fig. S2-S4).

### Literature review

We screened the Web of Science Core Collection database on December 28, 2020 using the following key words: “coronavir* AND (stability OR viability OR inactiv*) AND (temperature OR heat OR humidity)” (190 records). We also considered opportunistically found publications (23 records). We then selected the studies reporting original data focused on the effect of temperature on coronavirus inactivation obtained in experimental conditions (Fig. S1). For each selected study, we recorded information on virus, suspension medium, container, incubator, temperature and humidity (Table S1).

## Data accessibility

Compiled literature data as well as code and data to reproduce the Bayesian estimation results and corresponding figures are available on Github: https://github.com/dylanhmorris/heat-inactivation [68]

## Supplemental Material file list

Supplemental file 1 - Supplemental text, tables (Table S1) and figures (Figures S1-S4). PDF file

## Acknowledgments

We thank Linsey C. Marr for helpful discussions. This research was supported by the Intramural Research Program of the National Institute of Allergy and Infectious Diseases (NIAID), National Institutes of Health (NIH). JOL-S, AG, and DHM were supported by the Defense Advanced Research Projects Agency DARPA PREEMPT (D18AC00031), JOL-S and AG were supported by the UCLA AIDS Institute and Charity Treks, and JOL-S was supported by the U.S. National Science Foundation (DEB-1557022), the Strategic Environmental Research and Development Program (SERDP, RC-2635) of the U.S. Department of Defense. The content of the article does not necessarily reflect the position or the policy of the US government, and no official endorsement should be inferred.

## References

1. Ong, S. W. X. et al. Air, Surface Environmental, and Personal Protective Equipment Contamination by Severe Acute Respiratory Syndrome Coronavirus 2 (SARS-CoV-2) from a Symptomatic Patient. JAMA (2020).

2. Cai, J. et al. Indirect Virus Transmission in Cluster of COVID-19 Cases, Wenzhou, China, 2020. Emerging Infectious Diseases 26 (2020).

3. Kwon, K.-S. et al. Evidence of Long-Distance Droplet Transmission of SARS-CoV-2 by Direct Air Flow in a Restaurant in Korea. Journal of Korean medical science 35 (2020).

4. Azimi, P., Keshavarz, Z., Laurent, J. G. C., Stephens, B. & Allen, J. G. Mechanistic transmission modeling of COVID-19 on the Diamond Princess cruise ship demonstrates the importance of aerosol transmission. Proceedings of the National Academy of Sciences 118 (2021).

5. van Doremalen, N. et al. Aerosol and Surface Stability of SARS-CoV-2 as Compared with SARS-CoV-1. New England Journal of Medicine, NEJMc2004973 (2020).

6. Matson, M. J. et al. Effect of Environmental Conditions on SARS-CoV-2 Stability inHuman Nasal Mucus and Sputum. Emerging Infectious Diseases 26 (2020).

7. Pfaender, S. et al. Environmental Stability and Infectivity of Hepatitis C Virus (HCV) in Different Human Body Fluids. Frontiers in microbiology 9, 504 (2018).

8. Lopman, B. et al. Environmental Transmission of Norovirus Gastroenteritis. Current opinion in virology 2, 96–102 (2012).

9. Breban, R., Drake, J. M., Stallknecht, D. E. & Rohani, P. The Role of Environmental Transmission in Recurrent Avian Influenza Epidemics. PLoS computational biology 5, e1000346 (2009).

10. Rogers, W. in Sterilisation of Biomaterials and Medical Devices 20–55 (Elsevier, 2012).

11. Wigginton, K. R., Pecson, B. M., Sigstam, T., Bosshard, F. & Kohn, T. Virus Inactivation Mechanisms: Impact of Disinfectants on Virus Function and Structural Integrity. Environmental Science & Technology 46, 12069–12078 (2012).

12. Chang, L., Yan, Y. & Wang, L. Coronavirus disease 2019: coronaviruses and blood safety. Transfusion medicine reviews 34, 75–80 (2020).

13. Henwood, A. F. Coronavirus disinfection in histopathology. Journal of Histotechnology 43, 102–104 (2020).

14. Fischer, R. J. et al. Effectiveness of N95 Respirator Decontamination and Reuse againstSARS-CoV-2 Virus. Emerging Infectious Diseases 26, 2253 (2020).

15. Heimbuch, B. K. et al. A Pandemic Influenza Preparedness Study: Use of EnergeticMethods to Decontaminate Filtering Facepiece Respirators Contaminated with H1N1 Aerosols and Droplets. American Journal of Infection Control 39, 265–270 (2011).

16. Centers for Disease Control and Prevention. Implementing Filtering Facepiece Respirator (FFR) Reuse, Including Reuse after Decontamination, When There Are Known Shortages of N95 Respirators 2020.

17. Darnell, M. E. R. & Taylor, D. R. Evaluation of Inactivation Methods for Severe Acute Respiratory Syndrome Coronavirus in Noncellular Blood Products. Transfusion 46, 1770– 1777 (2006).

18. Leclercq, I., Batéjat, C., Burguière, A. M. & Manuguerra, J.-C. Heat Inactivation of the Middle East Respiratory Syndrome Coronavirus. Influenza and Other Respiratory Viruses 8 (2014).

19. Pagat, A.-M. et al. Evaluation of SARS-Coronavirus Decontamination Procedures. Applied Biosafety 12, 100–108 (2007).

20. Laude, H. Thermal Inactivation Studies of a Coronavirus, Transmissible GastroenteritisVirus. The Journal of General Virology 56, 235–240 (1981).

21. Pratelli, A. Canine Coronavirus Inactivation with Physical and Chemical Agents. TheVeterinary Journal 177, 71–79 (2008).

22. Duan, S.-M. et al. Stability of SARS Coronavirus in Human Specimens and Environment and Its Sensitivity to Heating and UV Irradiation. Biomedical and Environmental Sciences 16, 246–255 (2003).

23. Abu-Aisha, H. et al. The Effect of Chemical and Heat Disinfection of the Hemodialysis Machines on the Spread of Hepatitis c Virus Infection: A Prospective Study. Saudi Journal of Kidney Diseases and Transplantation 6, 174 (1995).

24. Jeong, E. K., Bae, J. E. & Kim, I. S. Inactivation of Influenza A Virus H1N1 byDisinfection Process. American Journal of Infection Control 38, 354–360 (2010).

25. Eterpi, M., McDonnell, G. & Thomas, V. Disinfection Efficacy against ParvovirusesCompared with Reference Viruses. Journal of Hospital Infection 73, 64–70 (2009).

26. World Health Organization. Guidelines on Sterilization and Disinfection Methods Effective against Human Immunodeficiency Virus (HIV) tech. rep. (1989).

27. Morris, D. H. et al. Mechanistic theory predicts the effects of temperature and humidity on inactivation of SARS-CoV-2 and other enveloped viruses. Elife 10, e65902 (2021).

28. Chin, A. W. H. et al. Stability of SARS-CoV-2 in Different Environmental Conditions. The Lancet Microbe 1, e10 (2020).

29. Yang, W. & Marr, L. C. Mechanisms by Which Ambient Humidity May Affect Viruses in Aerosols. Applied and Environmental Microbiology 78, 6781–6788 (2012).

30. Rockey, N. et al. Humidity and Deposition Solution Play a Critical Role in Virus Inactivation by Heat Treatment of N95 Respirators. Msphere 5 (2020).

31. Yang, W., Elankumaran, S. & Marr, L. C. Relationship between Humidity and Influenza a Viability in Droplets and Implications for Influenza’s Seasonality. PLoS ONE 7 (2012).

32. Benbough, J. E. Some Factors Affecting the Survival of Airborne Viruses. The Journal of General Virology 10, 209–220 (1971).

33. Lin, K., Schulte, C. R. & Marr, L. C. Survival of MS2 and ϕ6 viruses in droplets as a function of relative humidity, pH, and salt, protein, and surfactant concentrations. Plos one 15, e0243505 (2020).

34. Firquet, S. et al. Survival of enveloped and non-enveloped viruses on inanimate surfaces. Microbes and environments, ME14145 (2015).

35. Tuladhar, E., Bouwknegt, M., Zwietering, M., Koopmans, M. & Duizer, E. Thermal stability of structurally different viruses with proven or potential relevance to food safety. Journal of applied microbiology 112, 1050–1057 (2012).

36. Estep, T. N., Bechtel, M. K., Miller, T. J. & Bagdasarian, A. Virus Inactivation in Hemoglobin Solutions by Heat. Biomaterials, Artificial Cells and Artificial Organs 16, 129–134 (1988).

37. Gertsman, S. et al. Microwave-and Heat-Based Decontamination of N95 Filtering Facepiece Respirators: A Systematic Review. Open Science Framewor (2020).

38. Hulst, M. M. et al. Study on Inactivation of Porcine Epidemic Diarrhoea Virus, PorcineSapelovirus 1 and Adenovirus in the Production and Storage of Laboratory Spray-DriedPorcine Plasma. Journal of Applied Microbiology 126, 1931–1943 (2019).

39. Chan, K.-H. et al. Factors Affecting Stability and Infectivity of SARS-CoV-2. Journal of Hospital Infection 106, 226–231 (2020).

40. Darnell, M. E. R., Subbarao, K., Feinstone, S. M. & Taylor, D. R. Inactivation of the Coronavirus That Induces Severe Acute Respiratory Syndrome, SARS-CoV. Journal of Virological Methods 121, 85–91 (2004).

41. Gundy, P. M., Gerba, C. P. & Pepper, I. L. Survival of Coronaviruses in Water andWastewater. Food and Environmental Virology 1 (2009).

42. Kariwa, H., Fujii, N. & Takashima, K. Inactivation of SARS Coronavirus by Means of Povidone-Iodine, Physical Conditions and Chemical Reagents. Dermatology 212, 119–123 (2006).

43. Kim, Y.-I. et al. Development of Severe Acute Respiratory Syndrome Coronavirus 2 (SARS-CoV-2) Thermal Inactivation Method with Preservation of Diagnostic Sensitivity. Journal of Microbiology 58, 886–891 (2020).

44. Pujols, J. & Segalés, J. Survivability of Porcine Epidemic Diarrhea Virus (PEDV) inBovine Plasma Submitted to Spray Drying Processing and Held at Different Time by Temperature Storage Conditions. Veterinary microbiology 174, 427–432 (2014).

45. Saknimit, M., Inatsuki, I., Sugiyama, Y. & Yagami, K. Virucidal Efficacy of Physico-Chemical Treatments against Coronaviruses and Parvoviruses of Laboratory Animals. Experimental Animals 37, 341–345 (1988).

46. Chan, K. H. et al. The Effects of Temperature and Relative Humidity on the Viability of the SARS Coronavirus. Advances in Virology 2011, 1–7 (2011).

47. Thomas, P. R. et al. Evaluation of Time and Temperature Sufficient to Inactivate Porcine Epidemic Diarrhea Virus in Swine Feces on Metal Surfaces. Journal of SwineHealth and Production 23, 84 (2015).

48. Hofmann, M. & Wyler, R. Quantitation, Biological and Physicochemical Properties of Cell Culture-Adapted Porcine Epidemic Diarrhea Coronavirus (PEDV). Veterinary microbiology 20, 131–142 (1989).

49. Unger, S. et al. Holder Pasteurization of Donor Breast Milk Can Inactivate SARS-CoV-2. Canadian Medical Association Journal 192, E1657–E1661 (2020).

50. Batéjat, C., Grassin, Q., Manuguerra, J.-C. & Leclercq, I. Heat Inactivation of the Severe Acute Respiratory Syndrome Coronavirus 2. bioRxiv, 2020.05.01.067769 (2020).

51. van Doremalen, N., Bushmaker, T. & Munster, V. Stability of Middle East Respiratory Syndrome Coronavirus (MERS-CoV) under Different Environmental Conditions. Eurosurveillance 18, 20590 (2013).

52. Riddell, S., Goldie, S., Hill, A., Eagles, D. & Drew, T. W. The Effect of Temperature on Persistence of SARS-CoV-2 on Common Surfaces. Virology Journal 17 (2020).

53. Daeschler, S. C. et al. Effect of Moist Heat Reprocessing of N95 Respirators on SARS-CoV-2 Inactivation and Respirator Function. CMAJ 192, E1189–E1197 (2020).

54. Biryukov, J. et al. Increasing Temperature and Relative Humidity Accelerates Inactivation of SARS-CoV-2 on Surfaces. mSphere 5 (2020).

55. Mullis, L., Saif, L. J., Zhang, Y., Zhang, X. & Azevedo, M. S. Stability of Bovine Coronavirus on Lettuce Surfaces under Household Refrigeration Conditions. Food Microbiology 30, 180–186 (2012).

56. Guionie, O. et al. An Experimental Study of the Survival of Turkey Coronavirus atRoom Temperature and +4C. Avian Pathology 42, 248–252 (2013).

57. Casanova, L., Rutala, W. A., Weber, D. J. & Sobsey, M. D. Survival of SurrogateCoronaviruses in Water. Water Research 43, 1893–1898 (2009).

58. Casanova, L. M., Jeon, S., Rutala, W. A., Weber, D. J. & Sobsey, M. D. Effects of Air Temperature and Relative Humidity on Coronavirus Survival on Surfaces. Applied and Environmental Microbiology 76, 2712–2717 (2010).

59. Teytelman, L., Stoliartchouk, A., Kindler, L. & Hurwitz, B. L. Protocols. io: virtual communities for protocol development and discussion. PLoS Biology 14, e1002538 (2016).

60. Posada, J., Redrow, J. & Celik, I. A Mathematical Model for Predicting the Viability of Airborne Viruses. Journal of Virological Methods 164, 88–95 (2010).

61. Lofgren, E., Fefferman, N. H., Naumov, Y. N., Gorski, J. & Naumova, E. N. Influenza Seasonality: Underlying Causes and Modeling Theories. Journal of Virology 81, 5429– 5436 (2007).

62. Holshue, M. L. et al. First Case of 2019 Novel Coronavirus in the United States. New England Journal of Medicine 382, 929–936 (2020).

63. Tennant, B. J., Gaskell, R. M. & Gaskell, C. J. Studies on the Survival of Canine Coronavirus under Different Environmental Conditions. Veterinary Microbiology 42, 255–259 (1994).

64. Fischer, R. et al. Ebola virus stability on surfaces and in fluids in simulated outbreak environments. Emerging infectious diseases 21, 1243 (2015).

65. Brownie, C. et al. Estimating Viral Titres in Solutions with Low Viral Loads. Biologicals 39, 224–230 (2011).

66. Gelman, A. et al. Bayesian Data Analysis, Third Edition (CRC Press, 2013).

67. Carpenter, B. et al. Stan: A Probabilistic Programming Language. Journal of statistical software 76 (2017).

68. Gamble, A. et al. Data from “Heat-treated virus inactivation rate depends strongly on treatment procedure: illustration with SARS-CoV-2” https://github.com/dylanhmorris/ heat-inactivation.

69. Kay, M., Kola, T., Hullman, J. R. & Munson, S. A. When (ish) is my bus? user-centered visualizations of uncertainty in everyday, mobile predictive systems in Proceedings of the 2016 CHI Conference on Human Factors in Computing Systems (2016), 5092–5103.

