## Supplemental Material for "Heat-treated virus inactivation rate depends strongly on treatment procedure: illustration with SARS-CoV-2"

### 1 Literature review

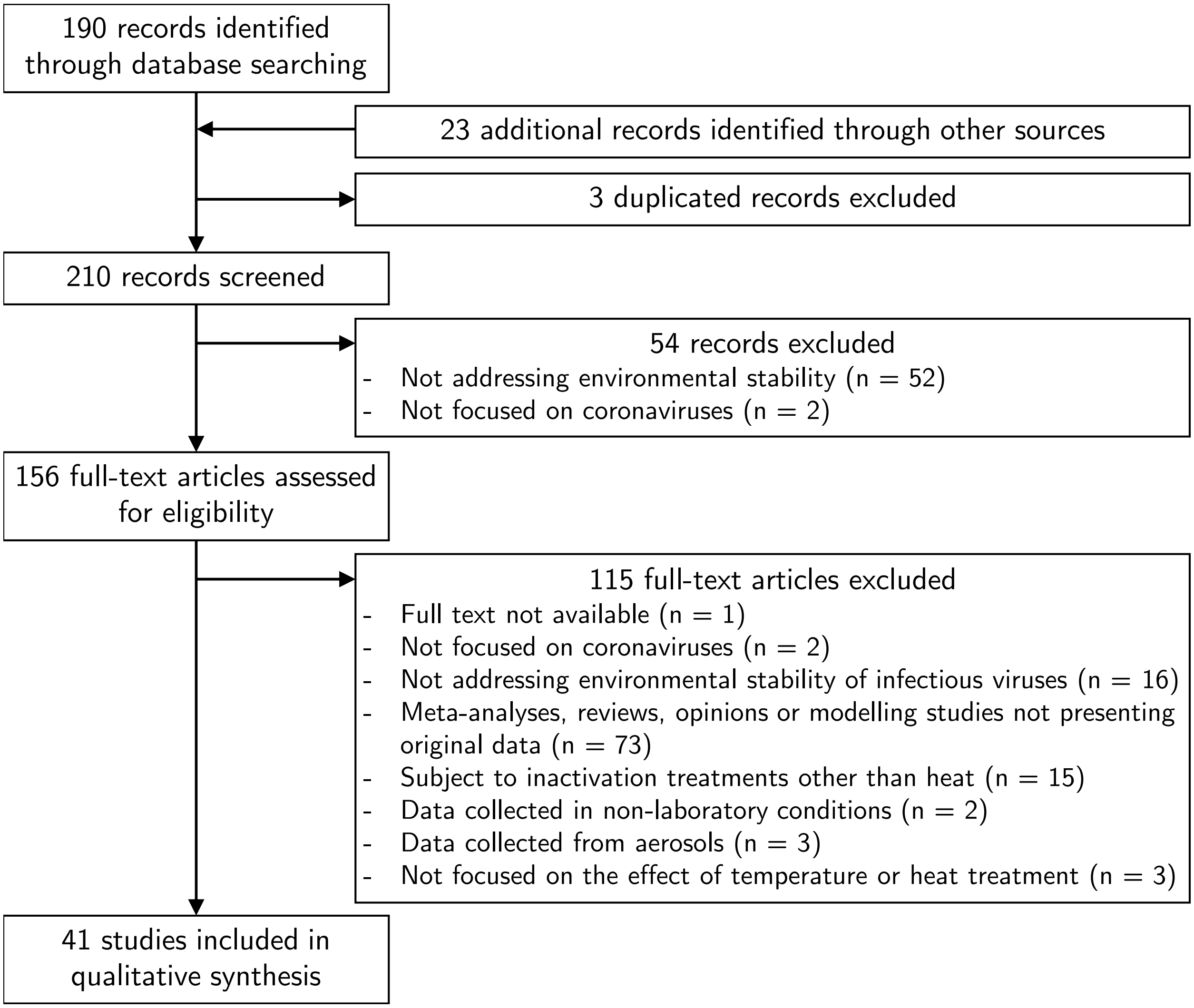

**Figure S1.** Selection process for literature review. Review assessed heat-treatment procedure description quality for coronavirus inactivation studies. This figure was made following the Preferred Reporting Items for Systematic Reviews and Meta-Analyses [1].

**Table S1.** Review of heat-treatment procedure description in the literature focusing on coronavirus inactivation. NI: not indicated; "?": information was not explicit. For instance Lai et al. 2005 [2] stated that "samples were incubated in closed containers", but it is not clear whether the "containers" refers to individual samples (e.g., vials) or batched of samples (e.g., a box aiming at limiting the variations of environmental conditions, as in Casanova et al. 2010[3] and Guionie et al. 2013 [4]). Information between parentheses were obtained from personal communications with the authors. Additional information (notably medium type, virus concentration, container material, and temperature and humidity conditions) is available online in the Github repository accompanying this manuscript [5].

| Study | Virus | Volume | Material | Container | Cover | Incubator |
|---|---|---|---|---|---|---|
| Batejat et al. 2020 [6] | SARS-CoV-2 | 500 µL | Bulk medium | NI | NI | Heat block |
| Biryukov et al. 2020 [7] | SARS-CoV-2 | 1, 5, 10 µL | Surface | NI | NI | Incubator |
| Bucknall et al. 1972 [8] | HCoV-229E, HCoV-OC43 | NI | Bulk medium | NI | NI | NI |
| Casanova et al. 2009 [9] | MHV, TGEV | 45 000 µL | Bulk medium | NI | NI | NI, refrigerator |
| Casanova et al. 2010 [3] | MHV, TGEV | 10 µL | Surface | NI | NI | Controlled-condition container |
| Chan et al. 2011 [10] | SARS-CoV-1 | 10 µL | Surface | Plate (24-well), vial | Cap, uncovered? | NI (nebulizer?) |
| Chan et al. 2020 [11] | SARS-CoV-2 | NI, 10, 300 µL | Bulk medium, surface | Vial | Cap, NI | NI |
| Chin et al. 2020 [12] | SARS-CoV-2 | NI | Bulk medium | NI (vial) | NI (cap) | NI |
| Christianson et al. 1989 [13] | FIPV, FIPV-TS | NI | Bulk medium | NI | NI | NI |
| Daeschler et al. 2020 [14] | SARS-CoV-2 | 5 µL | Surface | NI | NI | Incubator |
| Darnell et al. 2004 [15] | SARS-CoV-1 | 320 µL | Bulk medium | Vial (1.5 mL) | NI | Heat block |
| Darnell et al. 2006 [16] | SARS-CoV-1 | NI | Bulk medium | NI | NI | Water bath |
| Duan et al. 2003 [17] | SARS-CoV-1 | 100 µL | Surface | NI | NI | NI |
| Fischer et al. 2020 [18] | SARS-CoV-2 | 50 µL | Surface | NI (tray) | NI (uncovered) | NI (incubator), oven |
| Guionie et al. 2013 [4] | TCoV | 400 µL | Bulk medium | NI | NI | Cold room, isothermal box |
| Gundy et al. 2008 [19] | FIPV, HCoV-229E | 30 000 µL | Bulk medium | Vial | NI | NI |
| Harbourt et al. 2020 [20] | SARS-CoV-2 | 50 µL | Surface | NI | NI | NI |
| Hofmann et al. 1989 [21] | PEDV | 1000 µL | Bulk medium | NI | NI | Water bath |
| Hulst et al. 2019 [22] | PEDV | 2100 µL | Bulk medium | Vial | Cap | Water bath |
| Kariwa et al. 2006 [23] | SARS-CoV-1 | NI | Bulk medium | Vial (50 mL) | NI | Water bath |
| Kim et al. 2020 [24] | SARS-CoV-2 | 500 µL | Bulk medium | Vial | NI | Heat block |
| Lai et al. 2005 [2] | SARS-CoV-1 | 3000 µL | Bulk medium | NI | Closed? | NI |
| Lamarre et al. 1989 [25] | HCoV-229E | NI | Bulk medium | NI | NI | NI |
| Laude et al. 1981 [26] | TGEV | 1000 µL | Bulk medium | Vial | NI | Water bath |
| Leclercq et al. 2014 [27] | MERS-CoV | 500 µL | Bulk medium | NI | NI | NI |
| Matson et al. 2020 [28] | SARS-CoV-2 | 50 µL | Bulk medium, surface | NI (tray), vial | NI (uncovered), sealed | NI (incubator) |
| Mullis et al. 2012 [29] | BCoV | 100 µL | Surface | NI | NI | Refrigerator |
| Pagat et al. 2007 [30] | SARS-CoV-1 | NI | Bulk medium | NI | NI | Water bath |
| Pratelli et al. 2008 [31] | CCV | 500 µL | Bulk medium | Vial (15 mL) | NI | NI |
| Pujols et al. 2014 [32] | PEDV | 1 g | Bulk medium | Vial (0.5 cm inner diameter) | NI | Water bath |
| Quist-Rybachuk et al. 2015 [33] | PEDV | 100 µL | Bulk medium | NI | NI | NI |
| Rabenau et al. 2005 [34] | SARS-CoV-1 | NI | Bulk medium | NI | NI | NI |
| Riddell et al. 2020 [35] | SARS-CoV-2 | 10 µL | Surface | NI | NI | Incubator |
| Rockey et al. 2020 [36] | MHV | 50 µL | Surface | NI | NI | Incubator |
| Saknimit et al. 1988 [37] | CCV, MHV | 1000 µL | Bulk medium | Vial | NI | Water bath |
| Tennant et al. 1994 [38] | CCV | 1000 µL | Bulk medium | Vial? | NI | NI |
| Thomas et al. 2015 [39] | PEDV | 5000 µL | Surface | Tray | NI | Incubator |
| Unger et al. 2020 [40] | SARS-CoV-2 | 1000 µL | Bulk medium | NI | NI | NI, water bath |
| van Doremalen et al. 2013 [41] | MERS-CoV | 5 µL | Surface | NI (tray) | NI (uncovered) | Incubator |
| Ye et al. 2016 [42] | MHV | 30 000 µL | Bulk medium | NI | NI | NI |
| Yunoki et al. 2004 [43] | SARS-CoV-1 | NI | Bulk medium | NI | NI | NI |

#### 2 Bayesian estimation models

##### 2.1 Model notation

In the model notation that follows, the symbol $\sim$ denotes that a random variable is distributed according to a given distribution. Normal distributions are parametrized as:

$$\text{Normal}(\text{mean}, \text{standard deviation})$$

Positive-constrained normal distributions ("Half-Normal") are parametrized as:

$$\text{Half-Normal}(\text{mode}, \text{standard deviation})$$

##### 2.2 Titer inference

We inferred individual titers directly from titration well data using a Poisson single-hit model. We assigned a weakly informative Normal prior to the $\log_{10}$ titers $v_i$ ($v_i$ is the titer for sample $i$ measured in $\log_{10}\text{TCID}_{50}/0.1\text{mL}$, since wells were inoculated with 0.1mL):

$$v_i \sim \text{Normal}(2.5, 3) \tag{1}$$

We then modeled individual positive and negative wells for sample $i$ according to a Poisson single-hit model [44]. That is, for an undiluted inoculum, the number of virions that successfully infect cells within a given well is Poisson distributed with mean:

$$\ln(2)10^{v_i} \tag{2}$$

The value of the mean derives from the fact that our units are $\text{TCID}_{50}$; the probability of a positive well at $v_i = 0$, i.e. 1 $\text{TCID}_{50}$, is equal to $1 - e^{-\ln(2)\times 1} = 0.5$.

Let $Y_{idk}$ be a binary variable indicating whether the $k^{\text{th}}$ well at dilution factor $d$ (where $d$ expressed as $\log_{10}$ dilution factor) for sample $i$ was positive (so $Y_{idk} = 1$ if that well was positive and 0 if it was negative). Under a single-hit process, a well will be positive as long as at least one virion successfully infects a cell.

It follows from equation 2 that the conditional probability of observing $Y_{idk} = 1$ given a true underlying $\log_{10}$ titer $v_i$ and a dilution factor $d$ is given by:

$$\mathcal{L}(Y_{idk} = 1 \mid v_i) = 1 - e^{-\ln(2)\times 10^{(v_i - d)}} \tag{3}$$

This is simply the probability that a Poisson random variable with mean $\ln(2) \times 10^{(v_i-d)}$ is greater than 0. That mean is the expected number virions inoculated into the well; it derives from the fact that there are $v_i - d$ $\log_{10}\text{TCID}_{50}$ in the dilute sample. Similarly, the conditional probability of observing $Y_{idk} = 0$ given a true underlying $\log_{10}$ titer $v_i$ is:

$$\mathcal{L}(Y_{idk} = 0 \mid v_i) = e^{-\ln(2)\times 10^{(v_i - d)}} \tag{4}$$

which is the probability that the Poisson random with variable is equal to 0.

This gives us our likelihood function, assuming independence of outcomes across wells. Our inoculated doses were of volume 0.1 mL, so we incremented inferred titers by 1 to convert to units of $\log_{10}\text{TCID}_{50}/\text{mL}$.

#### 2.3 Virus inactivation regression

Duration of virus of detectability depends not only on environmental conditions and treatment method but also initial inoculum and sampling noise. We therefore estimated the exponential decay rates of viable virus (and thus virus half-lives) using a Bayesian regression analogous to that used in [18, 45]. This modeling approach allowed us to account for differences in initial inoculum levels across samples as well as other sources of experimental noise. The model yields estimates of posterior distributions of viral decay rates and half-lives in the various experimental conditions – that is, estimates of the range of plausible values for these parameters given our data, with an estimate of the overall uncertainty [46].

Our data consist of four different experimental conditions corresponding to four heat-treatment procedures, all at 70°C: (1) an uncovered plate of wells in a dry oven, (2) a covered plate in the oven, (3) a set of closed vials in the oven, and (4) set of closed vials in a heat block.

For each treatment, we took three samples per time point at multiple time-points.

We model each sample $j$ for experimental condition $i$ as starting with some true initial $\log_{10}$ titer: $v_{ij0}$. At the time $t_{ij}$ that it is sampled, it has titer $v_{ij}$.

We assume that viruses in experimental condition $i$ decay exponentially at a rate $\lambda_i$ over time. It follows that:

$$v_{ij} = v_{ij0} - \lambda_i t_{ij} \tag{5}$$

We use the direct-from-well data likelihood function described above, except that now instead of titers we estimate $\lambda_i$ under the assumptions that our observed well data $Y_{idk}$ reflect the titers $v_{ij}$.

We assume that each experiment $i$ has a mean initial $\log_{10}$ titer $\bar{v}_{i0}$. We model the individual initial titers $v_{ij0}$ as normally distributed about $\bar{v}_{i0}$ with an estimated, experiment-specific standard deviation $\sigma_i$:

$$v_{ij0} \sim \text{Normal}(\bar{v}_{i0}, \sigma_i) \tag{6}$$

#### 2.4 Regression prior distributions

We placed a Normal prior on the mean initial $\log_{10}$ titers $\bar{v}_{i0}$ that reflects the known inocula.

$$\bar{v}_{i0} \sim \text{Normal}(4.5, 0.5) \tag{7}$$

We placed a Half-Normal prior on the standard deviations $\sigma_i$ that allows for potentially large variation (1 log) variation about the experiment mean, as well as for less variation:

$$\sigma_i \sim \text{Half-Normal}(0, 0.25) \tag{8}$$

To encode prior information about virus inactivation rate in an interpretable way, we placed a Normal prior on the log half-lives $\ln(h_i)$, where $h_i = \frac{\ln(2)}{\lambda_i}$:

$$\ln(h_i) \sim \text{Normal}(\ln(0.5), 2) \tag{9}$$

This prior reflects that both of rapid virus inactivation and substantially slower inactivation are plausible a priori.

#### 2.5 Predictive checks

We assessed the appropriateness of prior distribution choices using prior predictive checks and assessed goodness of fit for the estimated model using posterior predictive checks. The resultant checks are shown below.

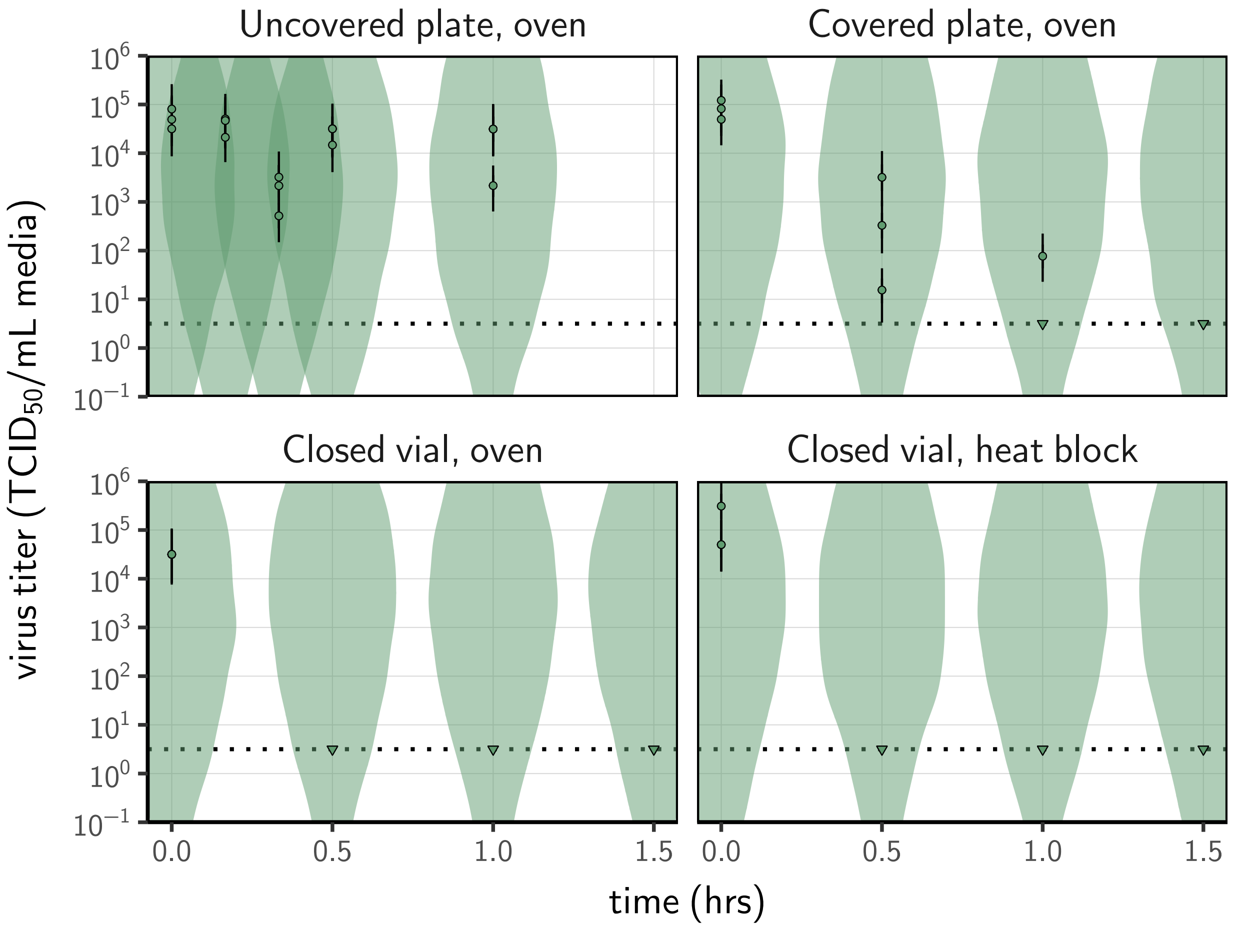

**Figure S2.** Titer estimation prior check. Violin plots show distribution of simulated titers sampled from the prior predictive distribution. Points show estimated titers for each collected sample; vertical bar shows a 95% credible interval. Time-points with no positive wells for any replicate are plotted as triangles at the approximate single-replicate detection limit of the assay (LOD; denoted by a black dotted line at $10^{0.5}$ $TCID_{50}$/mL media) to indicate that a range of sub-LOD values are plausible. Wide coverage of violins relative to datapoints show that priors are agnostic over the titer values of interest.

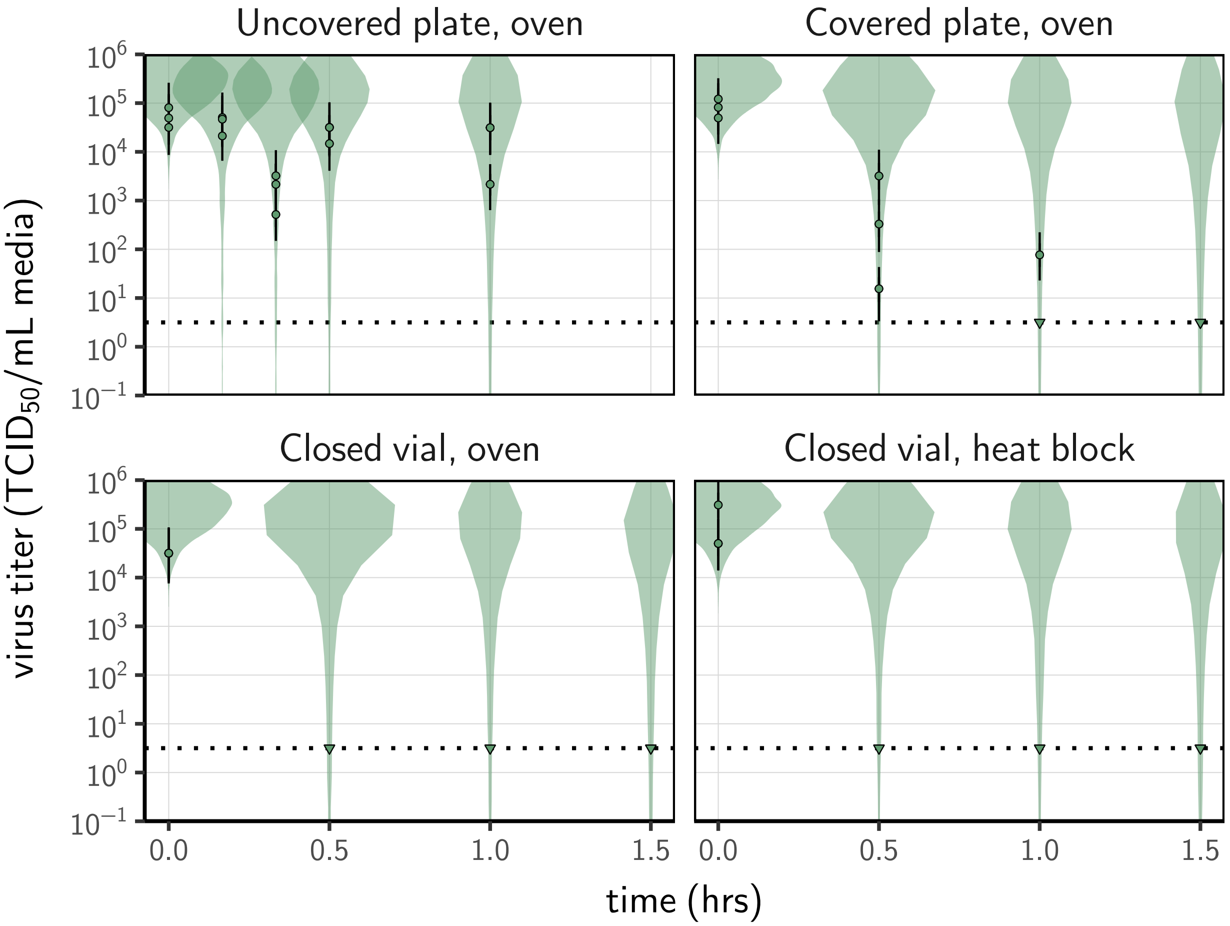

**Figure S3.** Prior predictive check for regression model. Violin plots show distribution of simulated titers sampled from the prior predictive distribution. Points show estimated titers for each collected sample; vertical bar shows a 95% credible interval. Time-points with no positive wells for any replicate are plotted as triangles at the approximate single-replicate detection LOD (denoted by a black dotted line at $10^{0.5}$ $TCID_{50}$/mL media) to indicate that a range of sub-LOD values are plausible. Wide coverage of violins relative to datapoints show that priors are agnostic over the titer values of interest, and that the priors regard both fast and slow decay rates as possible.

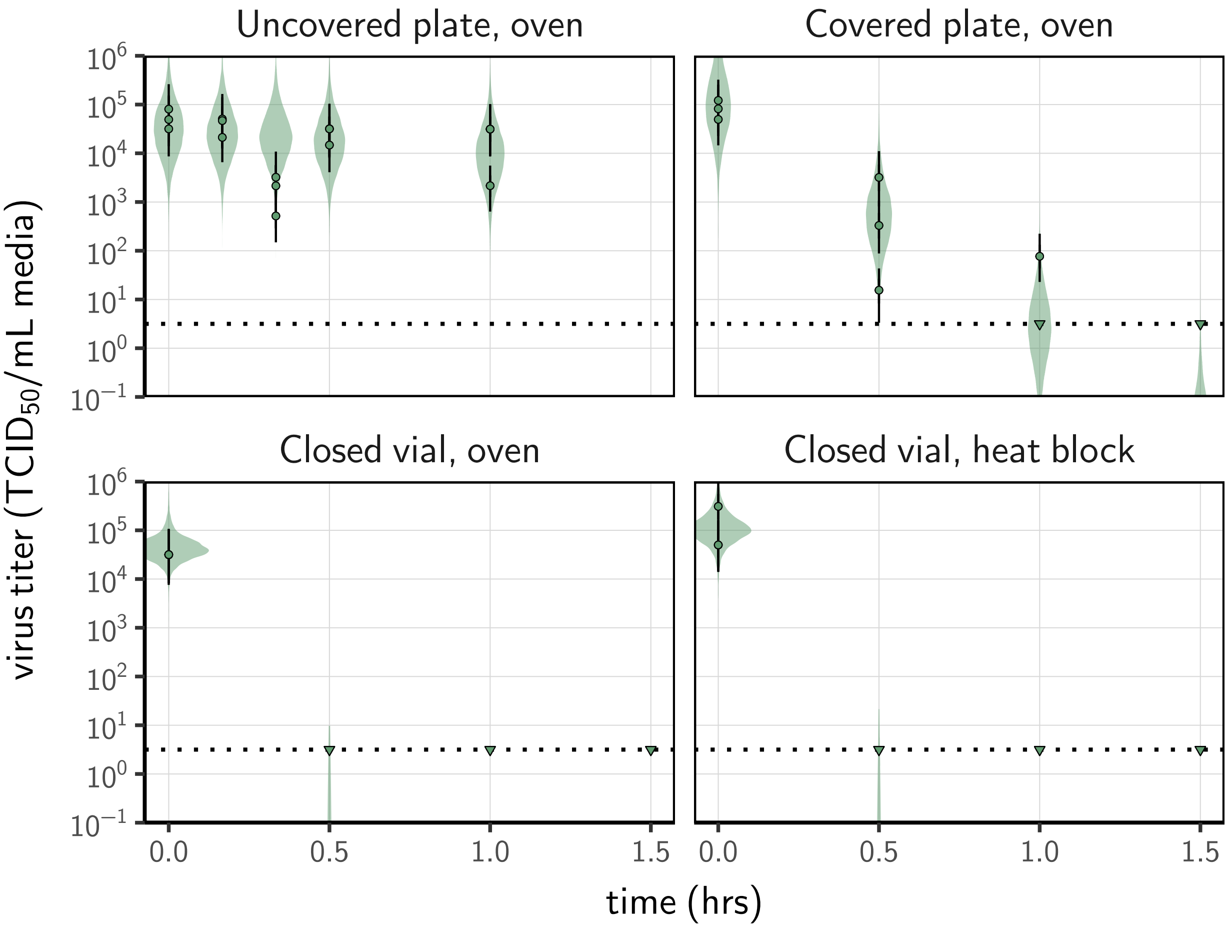

**Figure S4.** Posterior predictive check for regression model. Violin plots show distribution of simulated titers sampled from the posterior predictive distribution. Points show estimated titers for each collected sample; vertical bar shows a 95% credible interval. Time-points with no positive wells for any replicate are plotted as triangles at the approximate single-replicate detection LOD (denoted by a black dotted line at $10^{0.5}$ $\mathrm{TCID}_{50}$/mL media) to indicate that a range of sub-LOD values are plausible. Close correspondence between distribution of posterior simulated titers and estimated titers suggests the model fits the data well.

### References

1. Moher, D., Liberati, A., Tetzlaff, J. & Altman, D. G. Preferred Reporting Items for Systematic Reviews and Meta-Analyses: The PRISMA Statement. *PLoS Medicine* **6,** 6 (2009).
2. Lai, M. Y. Y., Cheng, P. K. C. & Lim, W. W. L. Survival of Severe Acute Respiratory Syndrome Coronavirus. *Clinical Infectious Diseases: An Official Publication of the Infectious Diseases Society of America* **41,** e67–71 (2005).
3. Casanova, L. M., Jeon, S., Rutala, W. A., Weber, D. J. & Sobsey, M. D. Effects of Air Temperature and Relative Humidity on Coronavirus Survival on Surfaces. *Applied and Environmental Microbiology* **76,** 2712–2717 (2010).
4. Guionie, O. *et al.* An Experimental Study of the Survival of Turkey Coronavirus at Room Temperature and +4C. *Avian Pathology* **42,** 248–252 (2013).
5. Gamble, A. *et al. Data from "Heat-treated virus inactivation rate depends strongly on treatment procedure: illustration with SARS-CoV-2"* https://github.com/dylanhmorris/heat-inactivation.
6. Batéjat, C., Grassin, Q., Manuguerra, J.-C. & Leclercq, I. Heat Inactivation of the Severe Acute Respiratory Syndrome Coronavirus 2. *bioRxiv,* 2020.05.01.067769 (2020).
7. Biryukov, J. *et al.* Increasing Temperature and Relative Humidity Accelerates Inactivation of SARS-CoV-2 on Surfaces. *mSphere* **5** (2020).
8. Bucknall, R. A., King, L. M., Kapikian, A. Z. & Chanock, R. M. Studies With Human Coronaviruses II. Some Properties of Strains 229E and OC43. *Proceedings of the Society for Experimental Biology and Medicine* **139,** 722–727 (1972).
9. Casanova, L., Rutala, W. A., Weber, D. J. & Sobsey, M. D. Survival of Surrogate Coronaviruses in Water. *Water Research* **43,** 1893–1898 (2009).
10. Chan, K. H. *et al.* The Effects of Temperature and Relative Humidity on the Viability of the SARS Coronavirus. *Advances in Virology* **2011,** 1–7 (2011).
11. Chan, K.-H. *et al.* Factors Affecting Stability and Infectivity of SARS-CoV-2. *Journal of Hospital Infection* **106,** 226–231 (2020).
12. Chin, A. W. H. *et al.* Stability of SARS-CoV-2 in Different Environmental Conditions. *The Lancet Microbe* **1,** e10 (2020).
13. Christianson, K., Ingersoll, J., Landon, R., Pfeiffer, N. & Gerber, J. Characterization of a Temperature Sensitive Feline Infectious Peritonitis Coronavirus. *Archives of virology* **109,** 185–196 (1989).
14. Daeschler, S. C. *et al.* Effect of Moist Heat Reprocessing of N95 Respirators on SARS-CoV-2 Inactivation and Respirator Function. *CMAJ* **192,** E1189–E1197 (2020).
15. Darnell, M. E. R., Subbarao, K., Feinstone, S. M. & Taylor, D. R. Inactivation of the Coronavirus That Induces Severe Acute Respiratory Syndrome, SARS-CoV. *Journal of Virological Methods* **121,** 85–91 (2004).
16. Darnell, M. E. R. & Taylor, D. R. Evaluation of Inactivation Methods for Severe Acute Respiratory Syndrome Coronavirus in Noncellular Blood Products. *Transfusion* **46,** 1770–1777 (2006).
17. Duan, S.-M. *et al.* Stability of SARS Coronavirus in Human Specimens and Environment and Its Sensitivity to Heating and UV Irradiation. *Biomedical and Environmental Sciences* **16,** 246–255 (2003).
18. Fischer, R. J. *et al.* Effectiveness of N95 Respirator Decontamination and Reuse against SARS-CoV-2 Virus. *Emerging Infectious Diseases* **26,** 2253 (2020).
19. Gundy, P. M., Gerba, C. P. & Pepper, I. L. Survival of Coronaviruses in Water and Wastewater. *Food and Environmental Virology* **1** (2009).
20. Harbourt, D. *et al.* Modeling the Stability of Severe Acute Respiratory Syndrome Coronavirus 2 (SARS-CoV-2) on Skin, Currency, and Clothing. *medRxiv,* 2020.07.01.20144253 (2020).
21. Hofmann, M. & Wyler, R. Quantitation, Biological and Physicochemical Properties of Cell Culture-Adapted Porcine Epidemic Diarrhea Coronavirus (PEDV). *Veterinary microbiology* **20,** 131–142 (1989).

22. Hulst, M. M. *et al.* Study on Inactivation of Porcine Epidemic Diarrhoea Virus, Porcine Sapelovirus 1 and Adenovirus in the Production and Storage of Laboratory Spray-Dried Porcine Plasma. *Journal of Applied Microbiology* **126,** 1931–1943 (2019).

23. Kariwa, H., Fujii, N. & Takashima, K. Inactivation of SARS Coronavirus by Means of Povidone-Iodine, Physical Conditions and Chemical Reagents. *Dermatology* **212,** 119–123 (2006).

24. Kim, Y.-I. *et al.* Development of Severe Acute Respiratory Syndrome Coronavirus 2 (SARS-CoV-2) Thermal Inactivation Method with Preservation of Diagnostic Sensitivity. *Journal of Microbiology* **58,** 886–891 (2020).

25. Lamarre, A. & Talbot, P. J. Effect of pH and Temperature on the Infectivity of Human Coronavirus 229E. *Canadian Journal of Microbiology* **35,** 972–974 (1989).

26. Laude, H. Thermal Inactivation Studies of a Coronavirus, Transmissible Gastroenteritis Virus. *The Journal of General Virology* **56,** 235–240 (1981).

27. Leclercq, I., Batéjat, C., Burguière, A. M. & Manuguerra, J.-C. Heat Inactivation of the Middle East Respiratory Syndrome Coronavirus. *Influenza and Other Respiratory Viruses* **8** (2014).

28. Matson, M. J. *et al.* Effect of Environmental Conditions on SARS-CoV-2 Stability in Human Nasal Mucus and Sputum. *Emerging Infectious Diseases* **26** (2020).

29. Mullis, L., Saif, L. J., Zhang, Y., Zhang, X. & Azevedo, M. S. Stability of Bovine Coronavirus on Lettuce Surfaces under Household Refrigeration Conditions. *Food Microbiology* **30,** 180–186 (2012).

30. Pagat, A.-M. *et al.* Evaluation of SARS-Coronavirus Decontamination Procedures. *Applied Biosafety* **12,** 100–108 (2007).

31. Pratelli, A. Canine Coronavirus Inactivation with Physical and Chemical Agents. *The Veterinary Journal* **177,** 71–79 (2008).

32. Pujols, J. & Segalés, J. Survivability of Porcine Epidemic Diarrhea Virus (PEDV) in Bovine Plasma Submitted to Spray Drying Processing and Held at Different Time by Temperature Storage Conditions. *Veterinary microbiology* **174,** 427–432 (2014).

33. Quist-Rybachuk, G. V., Nauwynck, H. J. & Kalmar, I. D. Sensitivity of Porcine Epidemic Diarrhea Virus (PEDV) to pH and Heat Treatment in the Presence or Absence of Porcine Plasma. *Veterinary Microbiology* **181,** 283–288 (2015).

34. Rabenau, H. F. *et al.* Stability and Inactivation of SARS Coronavirus. *Medical Microbiology and Immunology* **194,** 1–6 (2005).

35. Riddell, S., Goldie, S., Hill, A., Eagles, D. & Drew, T. W. The Effect of Temperature on Persistence of SARS-CoV-2 on Common Surfaces. *Virology Journal* **17** (2020).

36. Rockey, N. *et al.* Humidity and Deposition Solution Play a Critical Role in Virus Inactivation by Heat Treatment of N95 Respirators. *Msphere* **5** (2020).

37. Saknimit, M., Inatsuki, I., Sugiyama, Y. & Yagami, K. Virucidal Efficacy of Physico-Chemical Treatments against Coronaviruses and Parvoviruses of Laboratory Animals. *Experimental Animals* **37,** 341–345 (1988).

38. Tennant, B. J., Gaskell, R. M. & Gaskell, C. J. Studies on the Survival of Canine Coronavirus under Different Environmental Conditions. *Veterinary Microbiology* **42,** 255–259 (1994).

39. Thomas, P. R. *et al.* Evaluation of Time and Temperature Sufficient to Inactivate Porcine Epidemic Diarrhea Virus in Swine Feces on Metal Surfaces. *Journal of Swine Health and Production* **23,** 84 (2015).

40. Unger, S. *et al.* Holder Pasteurization of Donor Breast Milk Can Inactivate SARS-CoV-2. *Canadian Medical Association Journal* **192,** E1657–E1661 (2020).

41. van Doremalen, N., Bushmaker, T. & Munster, V. Stability of Middle East Respiratory Syndrome Coronavirus (MERS-CoV) under Different Environmental Conditions. *Eurosurveillance* **18,** 20590 (2013).

42. Ye, Y., Ellenberg, R. M., Graham, K. E. & Wigginton, K. R. Survivability, Partitioning, and Recovery of Enveloped Viruses in Untreated Municipal Wastewater. *Environmental science & technology* **50,** 5077–5085 (2016).

43. Yunoki, M., Urayama, T., Yamamoto, I., Abe, S. & Ikuta, K. Heat Sensitivity of a SARS-Associated Coronavirus Introduced into Plasma Products. *Vox Sanguinis* **87,** 302–303 (2004).

44. Brownie, C. *et al.* Estimating Viral Titres in Solutions with Low Viral Loads. *Biologicals* **39,** 224–230 (2011).

45. van Doremalen, N. *et al.* Aerosol and Surface Stability of SARS-CoV-2 as Compared with SARS-CoV-1. *New England Journal of Medicine,* NEJMc2004973 (2020).

46. Gelman, A. *et al.* *Bayesian Data Analysis, Third Edition* (CRC Press, 2013).
